# ebayGSEA: An improved Gene Set Enrichment Analysis method for Epigenome-Wide-Association Studies

**DOI:** 10.1101/454025

**Authors:** Danyue Dong, Tian Yuan, Shijie C. Zheng, Andrew E. Teschendorff

**Author notes:** Corresponding author Email: Andrew E Teschendorff.

## Abstract

**Motivation:** The biological interpretation of differentially methylated sites derived from Epigenome-Wide-Association Studies remains a significant challenge. Gene Set Enrichment Analysis (GSEA) is a general tool to help aid biological interpretation, yet its correct and unbiased implementation in the EWAS context is difficult due to the differential probe representation of Illumina Infinium DNA methylation beadchips.

**Results:** We present a novel GSEA method, called ebayGSEA, which ranks genes, not CpGs, according to the overall level of differential methylation, as assessed using all the probes mapping to the given gene. Applied on simulated and real EWAS data, we show how ebayGSEA may exhibit higher sensitivity and specificity than the current state-of-the-art, whilst also avoiding differential probe representation bias. Thus, ebayGSEA will be a useful additional tool to aid the interpretation of EWAS data.

**Availability and implementation:** *ebayGSEA* is available from *https://github.com/aet21/ebayGSEA*, and has been incorporated into the ChAMP Bioconductor package (*https://www.bioconductor.org*).

## Background

The number of Epigenome-Wide-Association Studies (EWAS) has grown rapidly, yet the biological interpretation of the differentially methylated sites found in these studies remains a significant problem [1,2]. EWAS typically use Illumina Infinium beadchips to measure DNA methylation (DNAm) at over 480,000 or 850,000 CpGs, depending on the beadchip version [3, 4], and genes represented on these chips may have widely different numbers of probes mapping to them [5]. It has been noted that this differential probe representation may cause significant bias when conducting differential methylation and GSEA, favouring genes with more probes [5, 6]. This is similar to the well-known bias of RNA-Seq differential expression calls towards longer genes, and for this reason, methods that adjust for this bias in RNA-Seq data have been adapted to the DNAm context [5]. However, drawing an analogy between RNA-Seq and DNAm data is also misleading, because in the RNA-Seq context the length of the gene only affects the reliability of the measured expression level, whereas in the DNAm context, the reliability of the measured DNAm level at a given CpG site does not depend on the number of probes mapping to the same gene. Thus, although genes with higher probe representation are more likely to be called differentially methylated, directly adapting methods from RNA-Seq data to DNAm beadchips may introduce other biases and still lead to suboptimal GSEA.

## Description

Here we present a novel GSEA method for Illumina DNAm data, with an empirical Bayes interpretation (thus called ebayGSEA), which overcomes the differential probe representation bias, whilst also avoiding some of the residual biases of current state-of-the-art methods like GSAmeth [5]. GSAmeth works by ranking differentially methylated CpGs (DMCs), selecting those that pass a genome-wide significance threshold, then mapping these to genes and finally to biological terms (pathways). Adjustment for differential probe representation is carried out at the gene-mapping stage, whereby the significance of the number of DMCs mapping to a given gene is assessed in relation to how many probes map to that gene. This, however, may result in two undesirable outcomes. First, genes in a pathway where a substantial fraction of marginal DMCs do not pass genome-wide significance levels may result in the enrichment of the pathway being missed (**Fig.1a-d, Supplementary Information**). Second, two pathways, matched for all variables (number of genes, probes mapping to each gene, number of genes containing at least 1 DMC) but differing widely in terms of the number or effect size of DMCs within a gene, will be ranked equally (**Supplementary Information**). ebayGSEA overcomes these problems by adapting the global test [7] to directly rank genes according to their overall level of differential methylation, as assessed using all of the probes that map to the given gene and in a manner which avoids favouring genes containing more probes. Subsequently, enrichment of biological terms is performed on this ranked list of genes using either a standard one-tailed Wilcoxon rank sum test (WT), or a recently introduced more powerful version known as the Known Population Median test (KPMT) [8]. As a result, in the first scenario considered above, affected genes will be relatively highly ranked via ebayGSEA (Fig.1c), and the ensuing ranked list leads to significant enrichment of the pathway (**Fig.1d**). In the second scenario, ebayGSEA will favour the pathway containing more DMCs, as required (**Supplementary Information**).

**Figure 1:**
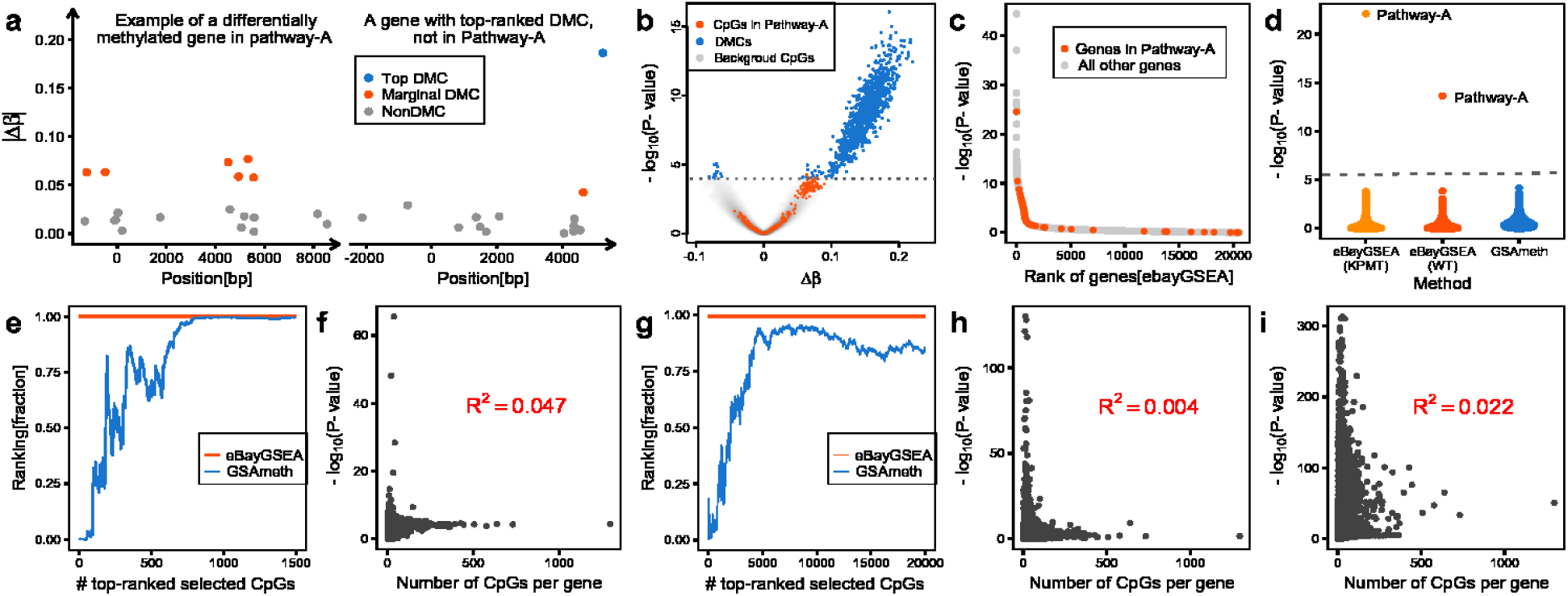
Validation of ebayGSEA. **a)** Example of a differentially methylated gene mapping to a hypothetical “pathway-A”, and of a gene containing a top-ranked DMC not mapping to pathway-A. y-axis labels the absolute differential methylation between two phenotypes. Each datapoint corresponds to a CpG mapping to the gene, with the position relative to the transcription start site (x=0). **b)** Volcano plot of the resulting DMCs with the grey dashed line indicating the line of significance (FDR=0.05). **c)** Significance (y-axis) vs. rank position of the gene (x-axis), as ranked by ebayGSEA. **d)** Significance of 8567 biological terms, as assessed using GSAmeth and ebayGSEA combined with either a Wilcoxon test (WT) or the Known Population Median test (KPMT). Dashed line marks Bonferroni threshold. **e)** Plot of the rank position (expressed as a fraction) of a biological term containing genes overexpressed in smoking related head&neck cancer in a smoking-EWAS performed in buccal swabs vs. the number of top-ranked selected CpGs used in GSAmeth (blue line). Red line indicates the rank position of the same term under ebayGSEA. **f)** Significance of the genes, as given by ebayGSEA, vs. the number of CpGs mapping to the gene, as derived using ebayGSEA in the same smoking EWAS. *R*^2^ value demonstrates that ebayGSEA is unbiased. **g-h)** As e-f, but now for a term of transcriptionally altered genes in an age-EWAS performed in blood. **i)** As panels f&h, but now for rheumatoid arthritis in an EWAS performed in blood.

We also compared ebayGSEA to GSAmeth in a smoking-EWAS performed on 400 buccal swabs [9]. Here, ebayGSEA ranked biological terms associated with smoking-related head&neck cancer much more highly than GSAmeth, the latter exhibiting wide variation depending on the number of top-ranked DMCs (Fig.1e, **Supplementary Information**). Of note, the ranking or statistical significance of genes derived from ebayGSEA did not correlate with the number of CpGs mapping to the gene, confirming that ebayGSEA, like GSAmeth, avoids differential probe representation bias (Fig.1f). Similar results were observed in other EWAS (Fig.1g-i,**Supplementary Information**).

## Conclusion

We propose that ebayGSEA be used alongside other GSEA methods to obtain a more objective and comprehensive assessment of GSEA in a given EWAS.

## References

1. Lappalainen T, Greally JM (2017) Associating cellular epigenetic models with human phenotypes. Nat Rev Genet 18:441–451.

2. Teschendorff AE, Relton CL (2018) Statistical and integrative system-level analysis of dna methylation data. Nat Rev Genet 19:129–147.

3. Beck S (2010) Taking the measure of the methylome. Nat Biotechnol 28:1026–8.

4. Moran S, Arribas C, Esteller M (2016) Validation of a dna methylation microarray for 850,000 cpg sites of the human. Epigenomics 8:389–99.

5. Phipson B, Maksimovic J, Oshlack A (2016) missmethyl: an r package for analyzing data from illumina’s humanmethylation450 platform. Bioinformatics 32:286–8.

6. Geeleher P, Hartnett L, Egan LJ, Golden A, l i Raja A, et al. (2013) Gene-set analysis is severely biased when applied to genome-wide methylation data. Bioinformatics 29:1851–7.

7. Goeman JJ, van de Geer SA, de Kort F, van Houwelingen HC (2004) A global test for groups of genes: testing an association with a clinical outcome. Bioinformatics 20:93–99.

8. Parks MM (2018) An exact test for comparing a fixed quantitative property between gene sets. Bioinformatics 34:971–977.

9. Teschendorff AE, Yang Z, Wong A, Pipinikas CP, Jiao Y, et al. (2015) Correlation of smoking-associated dna methylation changes in buccal cells with dna methylation changes in epithelial cancer. JAMA Oncol 1:476–85.

